# Brainwaves Monitoring via Human Midbrain Organoids Microphysiological Analysis Platform: MAP

**DOI:** 10.1101/2024.09.24.613225

**Authors:** SoonGweon Hong, Minsun Song, Woo Sub Yang, In-Hyun Park, Luke P. Lee

## Abstract

Understanding the development and pathogenesis of the human midbrain is critical for developing diagnostics and therapeutics for incurable neurological disorders including Parkinson’s disease (PD)^1–3^. While organoid models are introduced to delineate midbrain-related pathogenesis based on experimental flexibility^4–6^, there is currently a lack of tools with high fidelity for tracing the long-term dynamics of intact brain networks— an essential portrait of physiological states^7,8^. Here, we report a brain organoid microphysiological analysis platform (MAP) designed for long-term physiological development and in-situ real-time monitoring, akin to electroencephalogram (EEG), of midbrain organoids. We successfully achieved the on-chip homogeneous organogenesis of midbrain organoids and in-situ, non-disturbing electrophysiological tracking of the midbrain network activities. Throughout our long-term EEG monitoring via MAP, we captured the early-stage electrophysiological evolution of midbrain development, transitioning from discontinuous brief brainwave bursts to complex broadband activities. Furthermore, our midbrain organoid MAP facilitated the modeling and monitoring of neurotoxin-induced Parkinsonism, replicating the pathological dynamics of midbrain circuitry and exhibiting PD-like alterations in beta oscillation. We envision that the modeling and monitoring of brain organoid MAP will significantly enhance our understanding of human neurophysiology, neuropathogenesis, and drug discovery of neurodegenerative diseases.

## INTRODUCTION

Nestled within the uppermost segment of the brainstem lies the midbrain—a compact yet intricate region that assumes numerous essential physiological functions. As a central relay hub, this brain region plays a pivotal role in sensory processing, motor control, sleep regulation, and intricate reward and motivation. However, when neurological imbalances afflict this small domain, a spectrum of disorders emerges, encompassing conditions ranging from Parkinson’s disease to schizophrenia and addiction, unfolding over varying temporal scales.

The study of midbrain-associated neuropathology is a formidable task, hampered by the convoluted complexity of its anatomy, the intricate web of its functional dynamics, and its relatively modest size. While insights have been garnered from animal models and traditional cell cultures due to their easy accessibility, these non-human, non-physiological substitutes suffer from inherent discrepancies that temper the direct extrapolation of findings to human life sciences^9,10^. Recent advancements in comprehending the divergence in fetal brain development among species emphasize the need for more physiologically accurate representations of the human midbrain in research models^2,11^.

A recent human brain organoid modeling approach shows promise in mimicking how the midbrain develops and malfunctions^12–14^. Derived from human pluripotent stem cells, brain organoids undergo a three-dimensional self-organization mirroring the embryonic journey of the human brain^15^. Recent studies in single-cell RNA sequencing have unveiled the remarkable cellular fidelity of midbrain organoids to their in vivo counterparts, showing a symphony of cell types, including the crucial dopaminergic neurons meticulously programmed within the midbrain floor plate^16,17^. This ensemble extends to an array of midbrain non-dopaminergic populations—astrocytes, oligodendrocytes, GABAergic, glutamatergic, and serotonergic neurons—functional and flourishing within these organoids.

Researchers attempted to measure electrical activity in studies using these organoids to understand how the miniaturized brain models function^18^. Due to its lack of behavioral expression, scientists could learn by measuring how these cells’ electrical properties change, how the intercellular network communicates, reacts to stimuli, and changes during disease progression. This can help understand the causes of neurological disorders. Additionally, this method can be used to test new drugs, as it shows how compounds affect brain activity in a more lifelike setting^19^.

However, current methodologies exhibit certain limitations. For instance, patch clamping is an excellent technique for deciphering small-scale cellular processes. Yet, it falls short of elucidating how cells communicate within a larger network^20^. On the other hand, the multielectrode array (MEA) method offers insights into the spatial distribution of single-cell activity and computationally calculated network connections^21–23^ but can disrupt the tissue environment and architectures while establishing physical contact between neurons and electrodes, rendering it less than ideal for conducting extended longitudinal studies. Brain development and pathological progression are intricate processes that unfold over extended periods. These multifaceted events occur at various time scales, underscoring the need for a comprehensive and progressive examination to gain a deeper understanding of physiological dynamics.

This report introduces the midbrain organoid microphysiological analysis platform (MAP) (Fig. 1a). It is an integrated electrophysiological platform tailored to the cultivation of brain organoids and the in-situ exploration of their electrical symphony without disruption of the tissue environment. MAP facilitates the growing of multiple organoids in a uniform array with precise fluid dynamics (Fig. 1b), mimicking the brain’s development. Non-contact electrodes in the platform continuously monitor electrical activity without tissue disturbance (Fig. 1f vs. Fig. 1g), allowing us to track how midbrain networks develop, mature, and change over time (Fig. 1d). Using MAP, we recreated Parkinson’s disease-like symptoms in midbrain organoids by exposing them to a harmful substance. This allowed us to capture detailed electrophysiological progression that is otherwise challenging to observe, providing insights that complement the behavioral observations seen in animal models of Parkinsonism. We envisage that this platform will reshape our understanding of midbrain developmental trajectories and disease progression and could be valuable for testing drugs and personalized treatments for neurological disorders.

**Figure 1.**
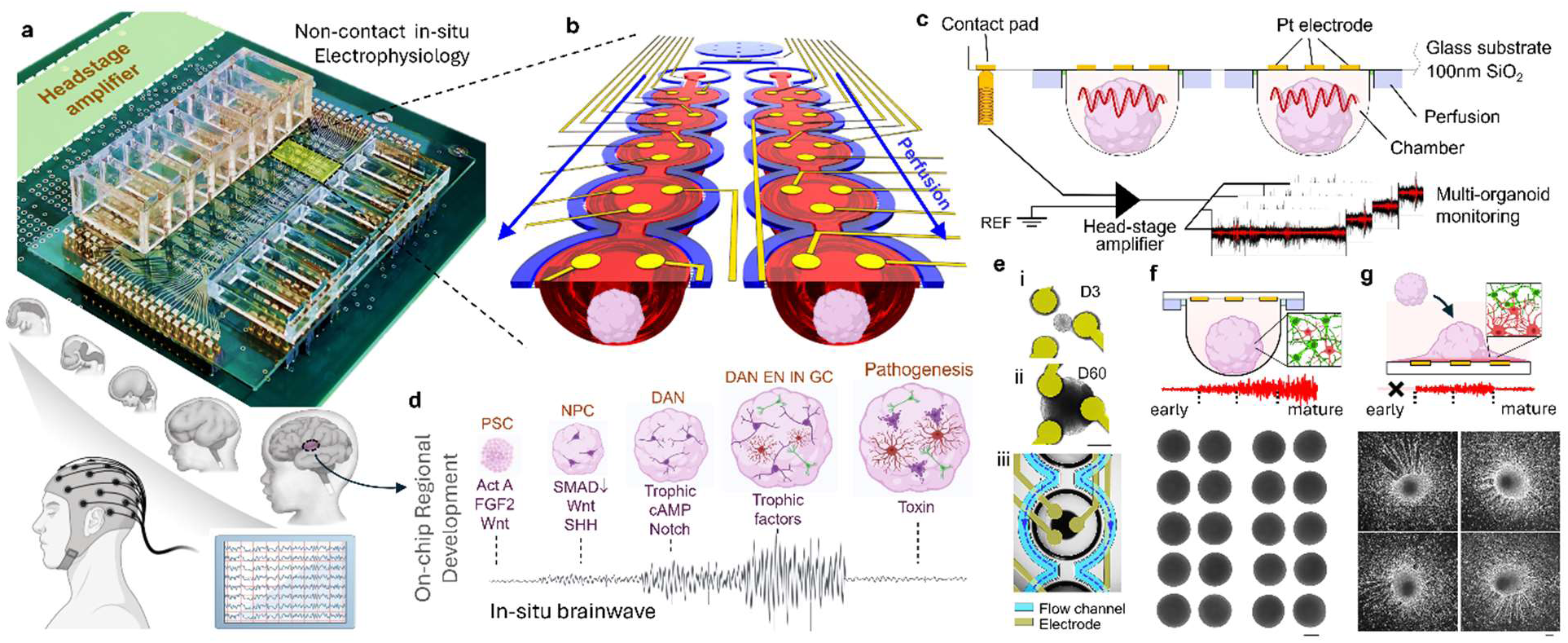
Microphysiological Analysis Platform (MAP) for In-Situ Electrophysiological Monitoring of Midbrain Organoids. **a,** A photograph of Brain Organoid MAP, comprising an integrated microfluidic device and a miniaturized electronic circuit for real-time electrophysiological monitoring of midbrain organoid development. The MAP layout allows for the generation of up to 80 brain organoids with dynamic perfusion facilitated by media reservoirs and a microprocessor-controlled tilting system. During on-chip organogenesis, the integrated electrical components sense extracellular potentials generated by brain organoids in a non-invasive, real-time manner, akin to clinical electroencephalogram (EEG) measurements. **b,** Close-up view of a functional unit within the MAP. Hemispherical organoid chambers (highlighted in red) are arrayed along perfusion channels (highlighted in blue). Each set of ten organoids is continuously exposed in a dynamic culture condition, surrounded by perfusion channels connecting an inlet and an outlet reservoir. The current MAP design includes eight biochemically independent units. Non-contact electrodes positioned above the hemispherical chambers enable EEG-like brainwave monitoring, exploiting the advantages of the hemispherical geometry for precise organoid positioning. **c,** Cross-sectional view of the integrated microfluidic device within the MAP, illustrating the multichannel electrical readout configuration. Surface electrodes, fabricated on a glass substrate through serial microfabrication involving platinum and silicon dioxide deposition, are composed of EEG electrodes on the top ceiling of organoid chambers, contact pads aligned to contact pins of a printed circuit board, and electric conduits under the dielectric layer. **d,** On-chip program for organogenesis, from a stem cell aggregate to a midbrain construct, followed by pathological modeling. Key biochemical components and their corresponding electrophysiological activities are indicated beneath schematics representing individual developmental stages, where PSC, NPC, DAN, EN, IN, and GC represent pluripotent stem cells, neural progenitor cells, dopaminergic neurons, excitatory neurons, inhibitory neurons, and glial cells, respectively. **e,** photographs of midbrain organoids in MAP. (i) & (ii) Top views of midbrain organoids and integrated electrodes during on-chip differentiation at day 3 and day 60, respectively. (iii) An expansive view with pseudocolors highlighting electrodes (yellow) and perfusion channels (blue with flow direction indicated by lines). **f & g,** Comparison of midbrain organoids cultured and monitored in MAP versus on MEA. On a MEA surface, the conformal contact of organoids leads to morphological changes, while MAP preserves the organoid architecture throughout the experiment, resulting in a more uniform morphology. Insets highlight astrocyte accumulation on the electrode surface of the MEA, whereas MAP maintains the innate brain cell distribution (with red and green cells representing astrocytes and neurons, respectively). The illustrative electrophysiological readouts demonstrate in-situ monitoring of the on-chip organogenesis in MAP. In contrast, MEA lacks early-stage readouts due to the use of a non-electrophysiological culture platform for initial organoid development and experiences poor signal quality because of astrocyte coverage. The organoid photographs illustrate the uniformity of midbrain organoids in MAP, as opposed to the deformation and uncontrollable architecture seen in midbrain organoids on MEA. Scale bar: 100 µm.

## RESULTS

### Microphysiological Analysis Platform (MAP) engineered for on-chip development of brain organoids and mapping brainwave dynamics by non-contact real-time EEG

Our microphysiological system^24^ provides an optimal microenvironment for maintaining metabolic homeostasis during electrophysiological activities and neurotransmitter synthesis of brain tissue. Through a microfluidic design resembling the body’s circulation (Fig. 1b), our MAP can continuously regulate the microenvironment by supplying nutrients without disrupting cellular mechanics or communication (Supplementary Fig. 1). This stability is crucial for long-term preserving the physiological integrity of brain organoids and ensuring their proper neuronal network development. Unlike traditional dish culture or flow methods that alter secretory signaling compared to in vivo conditions, MAP’s controlled perfusion around an interstitial space optimizes the regulation of differentiation factors during organoid formation. This yields reliable, consistent, reproducible brain organoid production, facilitating dependable quantitative analysis (Fig. 1f; Supplementary Fig. 2).

The microenvironment within the hemispherical chamber of MAP plays a pivotal role in generating and studying midbrain organoids. It not only maintains organoid positioning and aids visual tracking but also accommodates their growth from micrometer-scale assemblies to millimeter-sized structures (Fig. 1e). This scalability is essential for prolonged observations of developmental and pathological processes. Utilizing the hemispherical chamber, we’ve seamlessly integrated high-density surface electrodes positioned near organoids without direct contact. This configuration enables the capture of tissue-level electrical activity, distinct from conventional MEAs. The integration of electrical monitoring and microfluidic perfusion allows for simultaneous organoid culture and observation.

Our current platform is designed to support up to 80 organoids with parallel perfusion dynamics. It achieves eight independent biochemical modulations, while each organoid is monitored by three non-contact electrodes for electroencephalogram (EEG)-like readout, with reference electrodes exposed to the upstream fresh perfusion (Fig. 1c). The miniaturized system design allows for MAP integration with 256-channel head-stage amplifiers within a cell culture incubator. Additionally, our home-built tilting system regulates perfusion rates by controlling the angular position through preset calculations and integrated sensors.

To observe the full organotypic development of midbrain-specific structures, our initial approach involves creating an assembly analogous to embryoid bodies by introducing single-cell dissociated pluripotent stem cells into dedicated organoid chambers. The distinctive curvature of the hemispherical chamber is critical in rapidly re-aggregating dissociated stem cells, resulting in spheroids approximately 180 micrometers in diameter with a tight variation (Supplementary Fig. 2). Sequential application of differentiation protocols within the perfusion system (Fig. 1d) guides the stem cells through progressive stages—transforming into neuroectodermal cells, then into floor plate progenitors, and ultimately into midbrain-like structures that, over several months, exhibit spontaneous neuronal activity (Supplementary Fig. 5).

### MAP enables non-invasive, long-term, disturbance-free electrophysiology of organoid activities

Compared to conventional electrophysiological methods that involve direct cell-electrode contact, such as MEA systems, our MAP enables in-situ real-time monitoring alongside on-chip organogenesis while preserving organoid architecture (Fig. 2). Our proof-of-concept comparison highlights potential pitfalls associated with conventional electrophysiological approaches, particularly those resulting from neglecting the preservation of organoid architecture. To examine the consequences of organoid placement on an MEA surface, we compared the developmental variations of early-stage organoids between one group that transitioned to an MEA substrate at the floor-plate progenitor stage and a control group that remained intact MAP culture. This head-to-head comparison revealed the impact of midbrain organoid placement on MEA across four key aspects: morphological divergences, gene expression, cellular distribution, and electrophysiological readouts.

**Figure 2.**
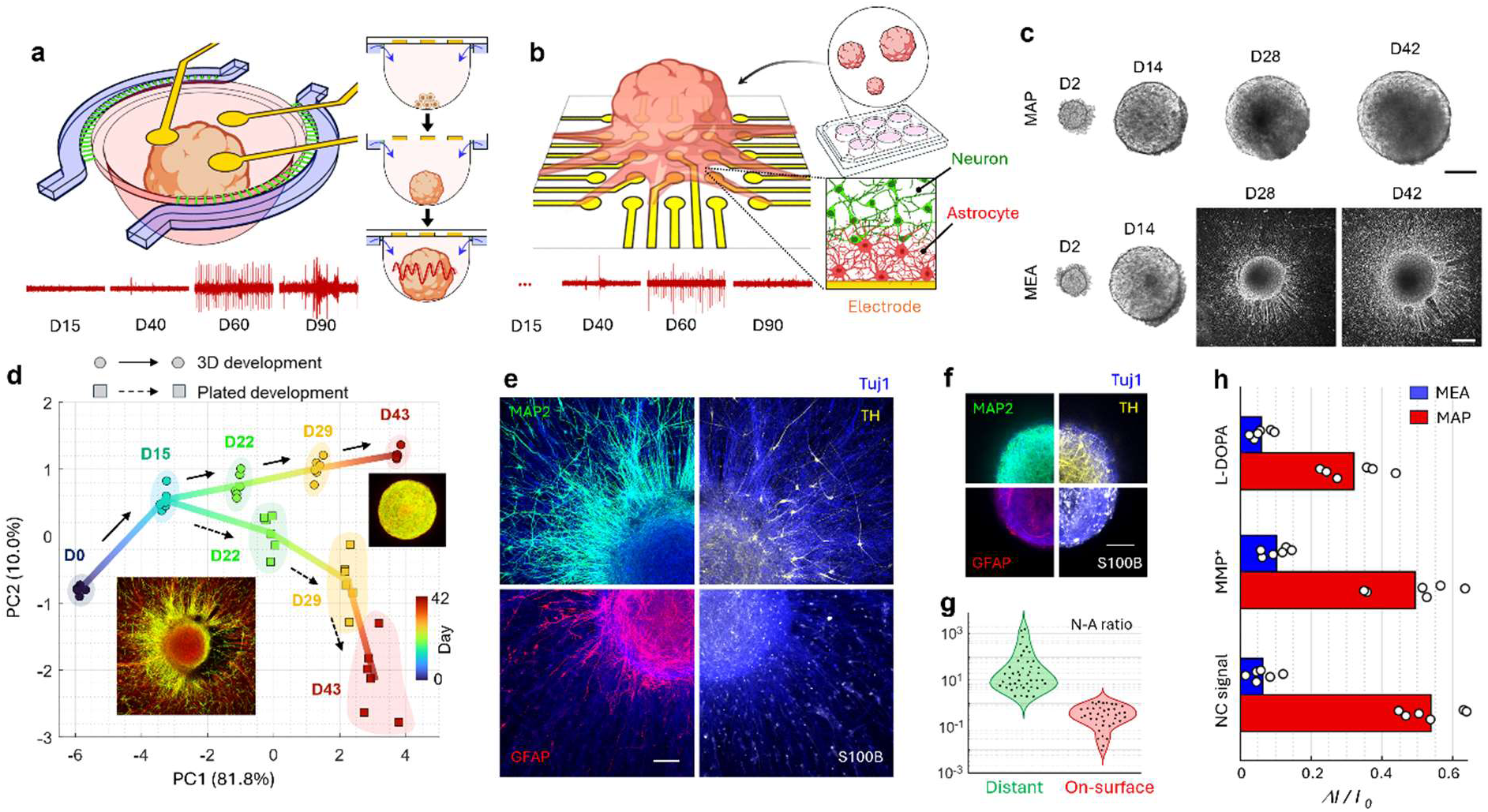
Comparative advantages of MAP midbrain organoid electrophysiology. **a & b,** The advanced capability of MAP in on-chip organogenesis and electrophysiology compared to the conventional MEA approach. The MAP’s direct formation of stem cell aggregation and on-chip differentiation facilitates in-situ electrophysiology of network activities of a brain construct without disturbing tissue architecture. In contrast, MEA experiments might overlook early-stage electrophysiological development because organoids are initially cultivated on a non-electrophysiological platform until they reach early stages. Additionally, these experiments may have less sensitive readouts of population activity due to abnormal astrocyte accumulation on the rigid surface. The time-lapse signals shown below each schematic represent the measurements from this comparative study. **c,** Time-trace photographs of midbrain organoids in MAP and MEA. The MEA organoid was cultured in MAP until the floor-plate progenitor stage (day 15) and then transferred to MEA coated with poly-l-ornithine and laminin. Scale bars of 200 µm are displayed in the day 42 MEA photograph for the plated images and are also shown in the day 42 MAP organoid images for the other images. **d,** Developmental trajectory characterized with midbrain-associated gene expression in MAP and MEA organoids. Both organoids were initially developed in the same MAP culture until the progenitor stage (day 15) to ensure consistency. One group continued in MAP for further development, while the other was transitioned to MEA. Representative immunostaining at Day 43 is shown in the insets, with MAP2 in green and Tuj1 in red. **e & f,** Distribution of MAP2, TH, GFAP, S100B, and Tuj1 across the MEA surface and within a section positioned one-quarter distance from the apex of the organoid. Scalar bars: 100 µm**. g,** Neuron-to-astrocyte ratio on the MEA surface and in the distant planes. **h,** Changes in MAP and MEA electrophysiological readouts on matured midbrain organoids in response to L-DOPA and MPP^+^, and NC (non-contact) signals. For L-DOPA and MPP^+^, the changes were calculated from time-averaged rectified LFP, while for the NC signal, it was calculated from the power spectrum at 15±1 Hz. The NC signal was externally applied as a 15 Hz sinusoidal oscillation to assess readout sensitivity to non-contact electrical signaling.

Upon placing the original spherical organoids on an MEA substrate, the organoids partially flattened to form conformal contact with the surface, as intended by the MEA protocol. The upper portion of the organoid, distant from the contact area, retained its three-dimensional morphology. However, as the post-progenitor development progressed, significant cellular outgrowth extended along the MEA surface from the contact area (Fig. 2c). This morphological deviation from the original three-dimensional spherical shape was linked to changes in the developmental trajectory of midbrain-associated mRNA expression as shown in Fig. 2d, with GFAP showing the highest upregulation, while most other genes were downregulated. The region of the organoid in contact with the MEA was rich in astrocytes but deficient in neurons, as indicated by GFAP and MAP2 expression (Figs. 2e, 2f & 2g). These changes likely had significant effects on the electrophysiological readouts, as MEA electrodes failed to capture the physiological contexts observed in the MAP record, leading to low-amplitude signals with poor signal-to-noise ratios in mature organoids on the MEA (Fig. 2a vs. Fig. 2b). Furthermore, the midbrain responses to L-DOPA and MPP^+^ did not yield statistically meaningful alterations in electrophysiological readouts from organoids on MEA (Fig. 2h), unlike the clear responses captured by MAP in the following sections. This underscores the limitations of MEA readouts in effectively examining the developmental physiology of brain organoids.

Our simulation models offer a potential rationale for the electrophysiological differences observed between MEA and MAP systems. Beyond the developmental deviations observed in the MEA-plated organoids, the accumulation of electrophysiologically less active astrocytes on the electrode surface disrupts the spatial propagation of electrical signals, hindering the transmission from more electrophysiologically active neurons situated farther from the electrode to the recording sensor (Fig. 4). This insensitivity to non-contact signals in the MEA configuration was confirmed by applying external signals through a proximally placed electrode pair (details in Methods), a limitation not observed in the MAP system (Fig. 2h). Furthermore, the MAP configuration enables additionally amplified signal readouts with the aid of the close chamber by minimizing potential dissipation across open boundaries (Supplementary Fig. 4), maintains a temporally consistent electrical background throughout the organoid study (Supplementary Fig. 8) and remains unaffected by fouling and degradation, as demonstrated in previous studies^25^. Thus, the non-contact electrophysiology provided by MAP enhances the long-term fidelity of electrophysiological studies of brain organoids and effectively captures population-oriented network activities while preserving organoid architecture and undisturbed development.

### MAP generates midbrain organoids rich in dopaminergic neurons’ presence and electrophysiological activity

Our MAP’s differentiation of stem cells to midbrain organoids allowed us to generate homogenous arrays of midbrain organoids whose overall development was well-matched to an anticipated trajectory with homogeneous expression levels across organoids (Fig. 3d & Fig. 2d). During differentiation initiation, pluripotency markers (NANOG & OCT4) swiftly diminished, indicating an initial commitment to specific lineages. In contrast, genetic markers (FOXA2, SOX1, NESTIN, LMX1B) tied to dopaminergic progenitors prominently emerged, becoming notably pronounced around day 15 of differentiation. Genes related to postmitotic neurons (MAP2, NEUN) and mature dopaminergic neuron development (TH, DAT, DDC, DRD2) gradually increased, mirroring trends observed in established models^26^. Among scrutinized genes, those linked to glial cell development (S100B, GFAP, AQP4) showed delayed development, consistent with prior studies^26^. Confocal microscopy of early-stage organoids revealed neurons with brief, slender MAP2-positive bodies and a modest presence of TH+ (dopaminergic) and S100B+ (glial) cells (Fig. 3a). Further maturation brought significant changes, including elongated neuronal extensions, an increased abundance of TH+ neurons, and the marked emergence of elongated glial cells. The confocal imaging additionally demonstrated minimal necrotic cores within the midbrain organoids, a phenomenon frequently encountered in brain organoid modeling ^27^. This may be attributed to our miniaturization of brain constructs facilitated by MAP and the continuous perfusion dynamics mimicking a physiological condition^28^ (Supplementary Fig. 1). Our single-organoid characterization demonstrates that the MAP’s differentiation promotes more consistent development of each organoid, both within and across batches, compared to other organoid generation methods.

**Figure 3.**
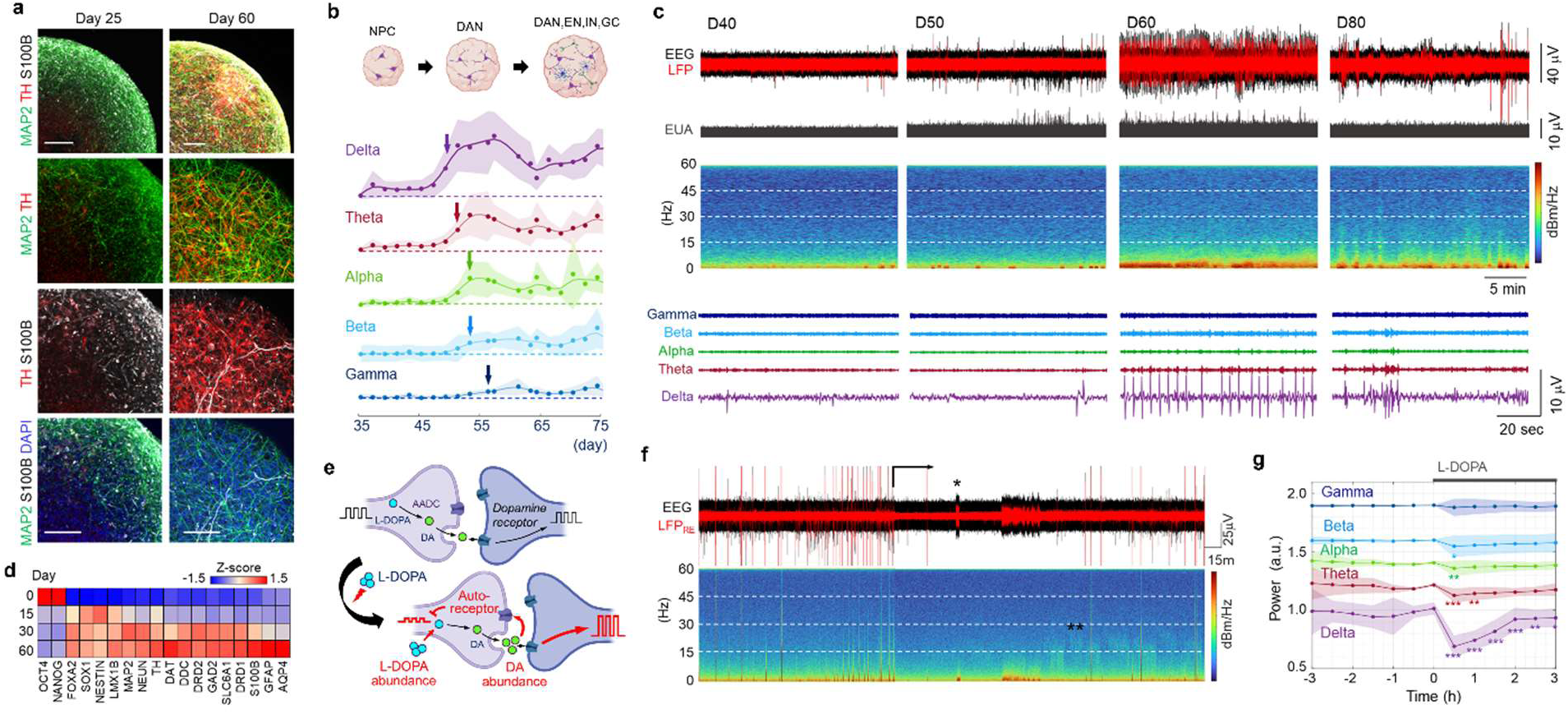
Characterization of Midbrain Organoids and Brainwave Evolution in MAP. **a,** Whole-mount immunostaining of midbrain organoids at day 25 and day 60, highlighting neuronal maturation (MAP2), dopaminergic neurons (TH), and glial cells (S100B). For visual clarity, DAPI is shown only in images with S100B and MAP2, presenting the distribution of cells in the organoid. These z-axis projections capture a quarter depth of the organoid size around a central plane. The matured organoid on day 60 exhibited well-defined MAP2^+^ neurons with a higher abundance of TH^+^ neurons and elongated glial morphologies compared to the earlier organoid. Scale bar: 100 μm. **b,** Temporal evolution of organoid-average brainwave powers during on-chip organoid differentiation. An accompanying schematic illustrates the expected electrophysiologically active components within developing midbrain organoids, where NPC, DAN, EN, IN, and GC represent neural progenitor cells, dopaminergic neurons, excitatory neurons, inhibitory neurons, and glial cells, respectively. Organoid-average brainwaves (n=10) are represented by lines with circle symbols, with filled areas indicating standard deviations. Arrows denote the earliest time points where significant deviations from baseline activities were observed, as determined by repeated measures ANOVA followed by Tukey’s Honestly Significant Difference (HSD) post-hoc analysis (*p* < 0.01). The analysis presents that midbrain organoids exhibit sequential turning points of brainwave development in a frequency-dependent manner. **c,** Electrophysiological readouts from MAP at differentiation days 40, 50, 60, and 80 reveal characteristic patterns throughout the electrophysiological development of midbrain organoids. The data, processed with a 60-Hz notch filter, are displayed as EEG. Local field potential (LFP) is shown as a 100 Hz low-pass signal, while unit spiking activity is represented by 300 Hz high-pass filtering and rectification. Spectrogram heatmaps on a logarithmic color scale illustrate the gradual transition from low-frequency-dominant to higher-frequency-containing brainwaves. Delta-to-gamma brainwaves displayed over a shorter period highlight characteristic episodes, showing the progression from brief bursts to continuous broadband waves. **d,** Transcriptomic profiles tracked during midbrain organoid differentiation in MAP. The colormap represents the relative expression of selected mRNAs using z-score normalization across the characterized time points. Pluripotency markers (OCT4, NANOG) rapidly decreased, while floor-plate progenitor markers (FOXA2, SOX1, NESTIN, LMX1B) increased at the beginning of differentiation. Genes related to postmitotic neurons (MAP2, NEUN) and dopaminergic maturation (TH, DAT, DDC, DRD2) demonstrated functional development for midbrain-specific networks. In contrast, genes associated with glial cells (S100B, GFAP, AQP4) developed at the later stages of differentiation. At each time point, gene expression was analyzed using three RNA pools from different batches, with each pool derived from ten organoids. **e,** Illustration depicting the electrophysiological response of midbrain organoids to L-DOPA treatment, centered on dopaminergic signaling. Presynaptic terminals indicate the effects on dopaminergic neurons synthesizing and releasing dopamine, while postsynaptic boutons represent neurons receiving enhanced dopamine signaling. Notably, dopaminergic signaling encompasses both synaptic and non-synaptic diffusive modes within the midbrain circuitry. The increased extracellular dopamine levels also trigger auto-receptor signaling in dopaminergic neurons. **f,** Electrophysiological response of a midbrain organoid (day 90) upon L-DOPA perfusion, initiated at the indicated point. LFP_RE_ corresponds to the local field potential obtained from 100 Hz lowpass filtering and a 20-millisecond time average for visualization. A post-L-DOPA burst train is indicated with *, while a transient, weak 10-30 Hz oscillation, not synchronized with the delta rhythm, is marked with **. **g,** Temporal changes in brainwave power upon L-DOPA treatment. Lines with circle symbols depict organoid-average power, while filled areas encompass the envelope with the standard deviation (*n*=20 from two batches). Statistical significance was assessed by performing a Student t-test against 20 age-matched vehicle control organoids (\**p* < 0.05, \*\**p* < 0.01, \*\*\**p* < 0.001).

To confirm that our organoid generated in MAP accurately models midbrain properties and has achieved functional maturity suitable for studying midbrain-specific electrophysiological responses, we investigated whether dopamine signaling manipulates midbrain organoid EEG expression using L-DOPA, a dopamine precursor to increase dopamine production in dopaminergic neurons (Fig. 3e) ^29,30^. Upon 50 µM L-DOPA application, the EEG power of midbrain organoids rapidly decreased, notably in the low-frequency region (Fig. 3f). Compared to age-matched vehicle control, the power decrease was significant in delta, theta, and alpha rhythm in a frequency-dependent manner while the significance became less over hours (Fig. 3g). A previous study suggests that the increase in extracellular dopamine induced by L-DOPA leads to decrease in dopaminergic neuron’s electrical firing via auto-receptor signaling, offering a potential explanation for the EEG power decrease^31^. Also, the midbrain organoids treated with L-DOPA exhibited sporadic emergence of electrophysiological episodes not observed before L-DOPA application or among the age-matched vehicle control organoids: elongated burst trains (noted with * in Fig. 3f) and fleeting occurrences of weak oscillations in the 10-30 Hz range as unsynchronized with the delta rhythm (noted in ** in Fig. 3f). An earlier study shows that dopamine-excited neurons in the midbrain slice respond to increased extracellular dopamine with higher firing frequencies than baseline activities of midbrain neurons^32^, which could potentially explain the secondary EEG episodes. While electrophysiological responses to L-DOPA stimulation reveal the activities of dopaminergic neurons central to EEG readouts of midbrain organoids, further investigations are crucial for gaining deeper insights into the secondary electrophysiological responses and the pharmacological effects of L-DOPA.

### MAP with integrated EEG maps brainwaves during midbrain organogenesis

An important advantage of brain organoid modeling in MAP is its capacity to illuminate the intricate neural circuitry during the early stages of brain electrophysiological development, providing an alternative approach to comprehending complex brain networks^33^. Anatomical challenges of accessing the midbrain region, especially during its nascent developmental phase, have impeded the study of midbrain electrophysiological development. In such circumstances, the electrophysiological tracing of midbrain organogenesis in MAP provides valuable insight into its developmental neurobiology.

The MAP with integrated EEG capability has effectively allowed us to track the trajectory of electrophysiological development during on-chip midbrain organogenesis. In the continuous EEG records, we have observed the gradual emergence of brainwave rhythms, commencing around day 40 of differentiation (Fig. 3b). Like other embryonic brain regions exhibiting spontaneous activities of low-frequency oscillations, our midbrain organoids initiated electrophysiological expression with the onset of low-frequency-bound (< 5Hz), discontinuous brainwave oscillations mainly emphasized in delta rhythm (herein termed as a local field potential (LFP) burst; Fig. 3c). These emerged briefly and infrequently as a biphasic waveform with extended inter-burst intervals, exhibiting a unique feature distinct from those observed in the cortical region.

Around 10 days further into development, there was a noticeable increase in the frequency of LFP bursts, accompanied by intensified spike activities, as depicted in Figure 3c. While the electrophysiological activities expressed in the local field potential continued to be bound by low frequencies, the EEG trace displayed various durations of increased spiking activities (SA), often asynchronous with LFP bursts. This SA-LFP temporal separation might reflect dopaminergic neuron physiology, particularly dopamine signaling distinct from classical synaptic transmission^34^.

Subsequent differentiation phases during organoid maturation led to the recurring appearance of high-magnitude LFP bursts, interspersed with secondary waves. These recurrent bursts occurred in elongated periodic events lasting over ten minutes, featuring gradually transient bursting rates between 0.1∼0.3 Hz. Quiescent periods often bridged the gaps between groups of recurrent bursts. As maturation progressed, the organoid EEG exhibited heightened complexity, marked by EEG patterns that defied concise classification into the earlier brief bursts. The EEG oscillation became nearly continuous with less distinct sub-unit patterns. Our analysis revealed a reduction in the amplitude of delta rhythms, while oscillations in frequencies beyond the delta range exhibited higher amplitudes than in earlier stages. These temporal courses can be understood as the developmental process of brain networks, evolving into extensive circuitries^35,36^ (Supplementary Fig. 5).

Biologically, organoids exhibited slightly varied EEG patterns during their evolution. Nonetheless, the average trace of organoid EEG patterns demonstrates sequential turning points for delta, theta, alpha, beta, and gamma rhythms between days 45 and 55 days of differentiation within MAP’s organogenesis (Fig. 3b).The initial emergence of slow-oscillating rhythm held precedence in EEG development, akin to the nascent brains’ development in vivo^37^, and the midbrain organoid circuitry at the MAP maturation stage exhibited brainwaves extending beyond the delta rhythm. Throughout our observation period, the midbrain organoid EEG showed the highest amplitude in the delta rhythm with gradually decreased amplitudes inversely proportional to its frequency in other brainwave bands. This phenomenon is frequently noted in neonatal brains^38^, possibly attributed to less mature circuitry when compared to their adult counterparts. Overall, the relationship between oscillation amplitude and frequency in the EEG signals followed a 1/f power law, a general feature of brain electrophysiology^39^. In parallel with efforts to characterize and define various types of intercellular communications linked to brainwave frequencies, normal brainwave features could serve as a reference for constructing pathological models.

### The EEG of midbrain organoids reveals the pathological dynamics induced by MPP^+^

In a maturation stage characterized by diverse brainwave patterns, our developed midbrain organoid served as a model to simulate midbrain-associated pathological features. The matured midbrain organoid prominently exhibited brainwaves central to dopaminergic signaling, as shown in the L-DOPA experiment, and demonstrated broad delta-to-gamma power spectral density, implying network connections with other neuronal subtypes (Fig. 3). To further explore its responses under pathological conditions, we could subject healthy midbrain organoids to a neurotoxin for the assessment of pathological responses of a midbrain construct. As a proof-of-concept study, we employed the dopaminergic neuron-specific neurotoxin, 1-methyl-4-phenylpyridinium (MPP^+^), which has been intensively studied with induced Parkinsonism. While various models, including humans, animals, organ slices, and organoids have shown MPP^+^-induced loss of dopaminergic neurons and acute PD-like symptoms^40–42^, electrophysiological characterization has not been extensively investigated. Hence, we aimed to correlate electrophysiological features seen in our EEG readouts with established molecular, physiological, and behavioral contexts.

Previous research has clarified that the toxicokinetics of MPP^+^ disrupts extracellular dopamine autoregulation, primarily due to its high affinity for the dopamine transporter (DAT). Furthermore, its accumulation on mitochondrial membranes disrupts the NADH dehydrogenase of mitochondrial Complex I, ultimately leading to cell death of dopaminergic neurons^43^ (Fig. 4a; Supplementary Fig. 7). To observe the electrophysiological manifestation of these intricate toxicokinetics, we introduced a low-dose MPP^+^ perfusion over two weeks while monitoring the organoid’s electrophysiological activities in MAP. By comparing the resulting electrophysiological dynamics with the baseline profile, we could analyze its pathological trajectory, shifting from normal baseline to hyperactivity and further to a state of electrical inactivity (Fig. 4b; Supplementary Fig. 9).

Upon initiation of 12-day perfusion with 10µM MPP^+^, the organoid’s EEG gradually transitioned into a hyperactive neuronal state, characterized by intensified alpha, beta, and gamma rhythms except delta rhythm, along with heightened spiking activity (Figs. 4b & 4e). This augmented broadband hyperactivity peaked between day 3 and 4 of MPP^+^ perfusion, subsequently diminishing. Throughout the initial 10-day perfusion, the EEG transformed from continuous to intermittent, marked by shorter and sporadic LFP bursts. The drastic reduction of EEG expression after the hyperactivity indicated the significant loss of network activities within the midbrain organoids due to MPP^+^ neurotoxicity. Our continuous monitoring of this gradual pathological progression enabled us to capture the successive electrophysiological dynamics, aligning with conventional understanding at their final stages, also revealing the transient electrophysiological traits that underscored the nuances of MPP^+^ toxicokinetics.

**Figure 4.**
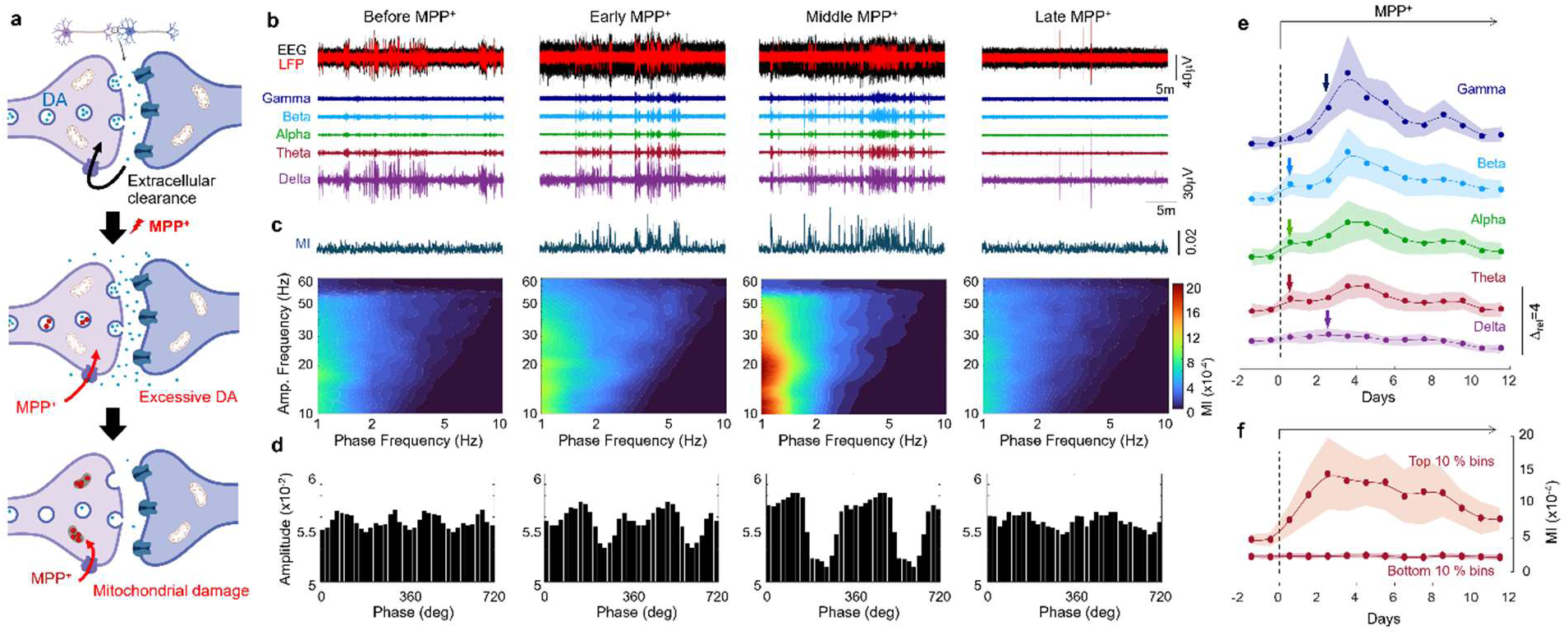
Brainwave Monitoring of MPP^+^-Induced Neurodegenerative Progress in Midbrain Organoids. **a,** Illustration outlining the toxicokinetic impact of MPP^+^ on midbrain dopaminergic signaling. The Dopamine Transporter (DAT) plays a pivotal role in regulating extracellular dopamine levels in midbrain tissue. MPP^+^’s affinity for DAT disrupts the extracellular clearance of dopamine, leading to abnormal excitation of dopamine-receiving populations. Internalized MPP^+^ within dopaminergic neurons progressively accumulates in mitochondria, disrupting the mitochondrial electron transport chain, and causing ATP depletion and eventual cell death. **b,** Distinct EEG and LFP patterns and corresponding brainwaves before and during chronic MPP^+^ application on day 90 midbrain organoids. The MPP^+^ stages, labeled as “Early,” “Middle,” and “Late,” represent 3, 8, and 12 days of MPP^+^ perfusion, respectively. The early and middle stages exhibit neuronal hyperactivity in the theta-to-gamma band, accompanied by increased spiking activity evident in EEG-LFP difference. The late stage is characterized by intermittent oscillations, marked by shorter and sporadic LFP bursts. **c,** Phase-amplitude coupling (PAC) analysis for brainwave alterations associated with MPP^+^ exposure, indicating abnormal delta-beta coupling during the MPP^+^-induced hyperactivity. Time-resolved traces of modulation index (MI) between the phase and amplitude frequency are computed at one-second intervals during the same time spans as the episodes in Panel b. During the hyperactivity, the MI levels change in synchronization with LFP bursts. Contoured heatmaps depict the time-average PAC comodulation for the presented episodes (span = 30 min). The later stage of hyperactivity (i.e., the middle MPP^+^) exhibits the highest level of delta-beta phase-amplitude coupling. **d,** Phase-amplitude coupling corresponding to the regions of maximal modulation index (MI) in the comodulation heatmap. The later hyperactivity period exhibits the most extensive coupling between delta and beta rhythms. **e,** Time-averaged brainwave power trends of midbrain organoids during MPP^+^ exposure. Organoid-average brainwaves (*n*=20 from two batches) are normalized against baseline values, with shaded areas representing standard deviations. The hyperactivity features peak between day 3 and 4 of MPP^+^ perfusion and gradually decrease to a state of electrophysiological inactivity. The arrows denote the earliest time points at which a significant difference emerged between MPP^+^-treated and age-matched vehicle control organoids (*n*=20 for each group) based on Student’s t-test analysis (*p* < 0.001). **f,** Temporal trace of MI values sorted by rectified LFP amplitude during MPP^+^ exposure. The time-resolved MI values are divided into the top and bottom 10% based on LFP amplitude for high and low electrophysiological activities. Organoid-average values (*n*=20 from two batches) are represented by lines with circle symbols, with shaded areas indicating standard deviations.

### Abnormal beta cross-couples to delta oscillation during MPP^+^-induced hyperactivity

Cross-frequency coupling (CFC) stands as a captivating phenomenon within neural oscillations. It involves the intricate interaction between brainwaves of varying frequencies, where the rhythm of one frequency is orchestrated by the rhythm of another. This intricate interplay between different frequency bands is considered to facilitate communication across disparate brain networks or regions. As such, the expression of CFC is a developmental and functional marker for the intricate circuitry that characterizes the brain’s operations^44–46^. Furthermore, disruptions or anomalies within the patterns of CFC hold the potential to serve as noteworthy indicators, shedding light on the underlying mechanisms that might contribute to an array of neurological and psychiatric disorders^47,48^.

In light of this, first, we examined the presence of CFC characteristics in a baseline matured state. To specifically identify CFC in midbrain organoid EEGs, we conducted an analysis that correlated the phase of delta rhythm with the amplitude of higher-frequency rhythms, achieved through Hilbert transform using a phase-amplitude coupling (PAC) analysis (details in Methods). The phase frequency range was between 1 and 10 Hz in 0.5 Hz resolution with a 1 Hz bandwidth, while the amplitude frequency ranged from 10 to 60 Hz in 1 Hz resolution with a 2 Hz bandwidth. This enabled us to establish a relationship between the central delta rhythm and the emerging higher-frequency oscillations. As illustrated by a representative time-averaged comodulation heatmap (Fig. 4c), matured midbrain organoids exhibited weak coupling between 1 Hz and a broad range of 10 to 50 Hz. The modulation index (MI), as observed in its time-resolved PAC trace and brainwaves, did not correlate with specific brainwave rhythms at this stage.

Upon introducing MPP^+^ perfusion, we observed a pronounced intensification of PAC of the delta rhythm particularly with the beta rhythm band (10-30 Hz). The delta-beta coupling peaked during the late phase of the hyperactivity, with a more distinct presence in the middle band (15-20 Hz) of the beta rhythm. Additionally, albeit relatively weak, a noticeable coupling between theta and low gamma frequencies persisted during the hyperactive state. As the EEG transitioned to an inactive state around the 12-day MPP^+^ treatment, the comodulation resembled that of the baseline EEG prior to the MPP^+^ application. The longitudinal comparison of PAC suggested that irregular delta-beta coupling was indicative of the hyperactive state (Figs. 4c & 4d).

The time-resolved MI trace demonstrated that delta-beta PAC exhibited a tendency to correlate with local field potential (LFP) bursts during the hyperactive phase (Fig. 4c). In contrast, this correlation was absent prior to MPP^+^ application, despite similar delta and beta rhythm amplitudes. Our analysis of daily MI distributions, sorted by rectified LFP amplitude to assess PAC dependency on electrophysiological activity, revealed that the PAC level during high LFP activities increased and peaked between day 2 and 3 of MPP^+^ perfusion (top 10% bins in Fig. 4f). These levels then gradually declined as the correlation tendency weakened, even though high MI values became more frequent during the later hyperactive stage. On the other hand, the PAC trace with reduced electrophysiological activity (i.e., bottom 10% bins in Fig. 4f) presented insignificant changes regardless of the MPP^+^ exposure. This suggests the presence of subtle, progressive electrophysiological changes during the MPP^+^-induced hyperactivity, which are not solely reflected in brainwave amplitude traces.

## DISCUSSION

This investigation presents a comprehensive microphysiological platform that facilitates the entire organogenesis of pluripotent stem cells into brain organoids within a physiological environment, and more importantly, enables real-time, long-term, and non-invasive electrophysiological monitoring of their intact developmental trajectory and pathological manifestations. Our dynamic culture environment supports extended physiological and pathological midbrain organoid progress, and its non-contact EEG-like readouts offer continuous monitoring of tissue-level electrophysiological dynamics over the extended period without imposing tissue disturbance. Through a four-month longitudinal study, we demonstrate the effectiveness of this approach in studying midbrain organoids, with indications that even longer culture and EEG recording periods are viable. The non-invasive nature of this platform is especially valuable when studying complex three-dimensional brain structures with tissue-level electrophysiological monitoring. Additionally, its compatibility with conventional analytical techniques such as immunostaining, microscopy, and transcriptomic analysis enhances its utility. The platform’s functional layout aligns with high-throughput assays in a standard multiwell format, with future scalability in mind. Amid the growing prominence of organoid research, our platform aims to improve developmental reproducibility, expedite research outcomes, and streamline organoid modeling.

In our demonstration of MEA electrophysiology on midbrain organoids, we highlight the critical concern that prolonged physical contact between electrodes and brain organoids causes adverse effects, especially in long-term electrophysiological studies. Our findings indicate that placing early-stage midbrain organoids on the MEA induces significant developmental deviations in both organoid transcriptomics and morphology, compared to architecture-preserving intact organoids. Similar to the biological response to chronically implanted electrodes in the brain^49^, we observed astrocyte accumulation on the MEA surface, which ultimately degraded signal quality despite the closer proximity to the organoid populations. These findings underscore the importance of more physiologically advanced electrophysiological techniques for organoids in progressive development, such as the non-contact, non-disruptive configuration provided by our MAP system.

Moreover, our MAP electrophysiology platform enables real-time, in-situ tracking of physiological activities throughout the entire organoid development process. In contrast, existing brain organoid electrophysiology methods typically require lengthy periods of differentiation in non-electrophysiological culture formats, followed by days to weeks of conformal contact with electrophysiological setups, and hours to days of cellular acclimation after media refreshment ^22,46,50–54^. These off-record periods inherently limit the ability to capture rare but crucial electrophysiological events. The early-stage brief discontinuous events, in particular, exemplify the importance of in-situ continuous monitoring.

Leveraging the in-situ monitoring capabilities of our platform during midbrain tissue development, this study uncovers unprecedented insights into the early developmental phase of the midbrain. During this phase, midbrain organoids exhibit brief, discontinuous activities that potentiate early neuronal network development. While such activities are common across various brain regions during fetal development, our study uniquely identifies midbrain-specific characteristics by comparing them with those in cortical organoids. Both organoid types exhibit brief LFP bursts characterized by large, slow, biphasic oscillations, with cortical EEG events generally showing higher amplitude. However, they differ significantly in the SA-LFP relationship. In cortical organoids, initial EEG events primarily synchronize with two temporally grouped bursts observed in entire unit activity (EUA; a mathematical expression of SA described in Methods), which are attributed to giant depolarizing potentials driven by early-stage GABA signaling^55^. In contrast, midbrain organoids show asynchrony between EUA and LFP bursts, suggesting distinct mechanisms of microcircuit development in nascent midbrain circuitry^56^. This observation underscores the diverse electrophysiological developmental origins across brain regions ^57,58^, which can be effectively studied through region-specific brain organoid models. While further studies will be needed to deepen the significance of these findings, the platform’s capability to trace intact electrophysiological trajectories during development presents a promising avenue for understanding neural circuit formation.

Another compelling aspect of our study is the observation of pathological dynamics induced in midbrain organoids by chronic MPP^+^ exposure. Characterizing electrophysiological properties at the level of specific organs has been historically limited due to the challenges of in vivo access, brain slice instability, and the complexity of body-wide interconnections. While the organoid model may not perfectly replicate in vivo biology with its long-term pathogenesis developed through body-wide connections, it offers a promising pseudo-ideal model specifying the brain region. The observed electrophysiological dynamics triggered by MPP^+^ application, transitioning from the transient hyperactivity to the electrophysiological inactive state, align with prior molecular findings. Previous studies have shown increased extracellular dopamine and glutamate in the midbrain following MPP^+^ application^59,60^. Independent research has demonstrated an increased firing rate of non-dopaminergic neurons and a decreased firing rate of dopaminergic neurons in response to MPP^+^ application^41^, supporting the notion of hyperactivity during MPP^+^ exposure. The observed delta-beta PAC may also reflect real-world phenomena, given that this pattern is commonly associated with anxiety^61,62^, a symptom observed in animal behavior studies after MPP^+^ injection and now linked as a potential precursor to Parkinson’s disease^63^. Although further validation through in vivo electrophysiological measurements in animal models is required, these insights suggest transient electrophysiological activities across multiple aspects of midbrain biology, increasing values of such miniaturized brain construct for close-up examination of pathological contexts.

From our perspective, the integrated MAP is poised to revolutionize the study of brain organoids, offering a means to comprehensively investigate neurogenesis, neuropathogenesis, and precise drug discovery. The MAP-enabled brain study allows us to (1) enable the entire process of on-chip organogenesis, from stem cells to uniform organoids, by precise controlling biophysical fluid dynamics to recapitulate physiological microenvironments; (2) monitor intact electrophysiological activity from the early embryonic development, shedding light on the fundamental processes of brainwave generation; and (3) intentionally perturbate physiological perfusion to induce controlled pathological conditions, facilitating a comprehensive exploration of neuropathogenesis through real-time brainwave recording. Besides, it is expected to revolutionize drug discovery and patient care by offering precision and personalized medicine for neurodegenerative diseases. This current study serves as an initial demonstration of MAP’s potential, harnessing an in-vitro brain organoid model to study the otherwise challenging-to-access brain region. Furthermore, common concerns associated with brain organoid models, such as standardization, lack of vascularization, scalability, and functional assessment, may be addressed and improved using the integrated MAP^64^. Our platform also has the potential to provide valuable insights into organoid maturity by minimizing unintended tissue stress^65^ that can expedite cellular aging in a developmentally uncorrelated manner. The anticipated broad impact of organoid MAP studies is on the horizon.

## METHODS

### Integrated MAP with Non-invasive Electrophysiology

The integrated microfluidic device (Fig. 1a) was constructed from three distinct components, each fabricated separately and subsequently assembled. The initial step involved creating the microfluidic unit using photolithography to generate a mold, followed by using polydimethylsiloxane (PDMS, DOW SYLGARD™ 184) for the polymer replication. While the procedure adhered closely to the methodology outlined in previous literature^24^, minor adjustments were made in the size of the organoid chamber for optimal organoid growth.

Simultaneously, the EEG electrodes were accurately patterned onto a glass slide (12.5cm x 6.5cm) utilizing a lift-off process. Adhering to the guidelines recommended by the manufacturer (Kayaku Advanced Materials), thin layers of LOR3A and SU8 3005 were initially patterned before the deposition of metals. This deposition involved the sequential application of a 5nm titanium layer and a 50nm platinum layer using an electron-beam evaporator (ROCKY Mountain Vacuum Tech). After a first lift-off to define the metal conduits using AZ 300 MIF (EMD), a similar process was employed to deposit a 100nm SiO_2_ layer as a passivation layer. Then, holes were carefully drilled on the electrode-patterned glass slide to enable fluid flow between the media reservoirs and the PDMS chip.

The third constituent, the media reservoirs, was fabricated by CO_2_ laser machining of 0.5” thick acrylic plates (VersaLaser VLS3.5). After laser machining, a thorough cleaning process was executed to ensure the removal of any residual debris. The final assembly phase involved joining all components through O_2_ plasma bonding, which secured the glass and the PDMS chip, and uncured PDMS bonding, which effectively connected the glass and the reservoirs.

The printed circuit board (PCB) for interfacing the MAP EEG electrodes was designed using CAD software and fabricated by a PCB manufacturer (Advanced Circuits, Inc.). Spring-loaded connectors (Mill-Max Manufacturing Corp.) were integrated into the PCB for physical connection to the contact pads of surface electrodes. Headstage amplifiers (RHD 64CH, Intan Technologies) were also integrated into the PCB for proximity to the recording sites. With the integration of the four amplifiers on the board, this board enabled simultaneous recording of 240 channels of MAP EEG at a recording rate of up to 20 kHz.

We carefully accounted for all components and layouts during the design phase to enable recording directly within a tissue culture incubator. After assembly, the combined dimensions of the MAP and PCB measured 20 cm by 40 cm. To reduce any interference from external noise sources, we devised a Faraday cage, using an aluminum enclosure to accommodate the entire assembly. We used a wired interface board from Intan Technologies, connected to a laptop for data acquisition. All data presented in the study was captured at a consistent recording rate of 5 kHz. The overall integration configuration can be referred to in our previous work^52^.

To ensure uninterrupted perfusion during recording, we devised a tilting plate system. This system seamlessly integrated a microcontroller (Arduino Uno, Adafruit Industries LLC), a position sensor (ADXL326, Adafruit Industries LLC), and a linear actuator motor (Pololu Corp.). The structural design comprised T-slotted framing rails, forming a spacious plane measuring 40 cm by 40 cm. This configuration was tailored to perfectly fit within a cell culture incubator and executed tilting motions following predetermined parameters. The tilting mechanism functioned on a continuous 24-hour cycle, with a daily reset synchronized to the moment of reservoir refilling. This approach ensured a consistent, controlled perfusion environment throughout the recording process without complicated tubing and pumping.

The MAP’s integration within the microfluidic configuration yields a noise level range of less than 35 µVpp with excellent temporal stability (Supplementary Fig. 8), demonstrating its adequacy for characterizing brainwave oscillations in this study. Furthermore, the temporal resolution, primarily determined by the headstage operation capable of reaching up to 30 kHz, has been deliberately set at 0.2 msec (equivalent to a 5 kHz readout) for this study, proving sufficient for our brainwave analysis.

### On-chip Organogenesis of Midbrain Organoids

We achieved the development of organoids within MAP, harnessing fluidic dynamics. The organoid differentiation process involved continuous delivery of differentiation factors through media perfusion, simulating in vivo floor plate signaling.

Our protocol began with cultivating human pluripotent stem cells (H9) in a Matrigel-coated 6-well plate using a stem cell feeder-free culture medium (mTeSR1, STEMCELL Technologies). For routine passaging, cell aggregates were gently detached using Gentle Cell Dissociation Reagent (GCDR, STEMCELL Technologies). To prevent spontaneous differentiation, regular scraping coupled with visual assessment was employed.

For loading onto the device, stem cells at 60-80% confluency were dissociated into single cells using Accutase (STEMCELL Technologies), followed by centrifugation and resuspension at a density of 0.3 million cells per milliliter in mTeSR1 supplemented with 10µM Y-27632 (a ROCK inhibitor). Before cell loading, the device was filled with PBS using a vacuum chamber to ensure the absence of air, which was confirmed under a microscope to prevent the introduction of electrical noise. The loaded cells were introduced into surface-passivated MAP organoid chambers in a bottom-up configuration. Following the sealing of loading inlets, the MAP device was flipped to its normal position, enabling cell aggregation on the base of the hemispherical chambers. An overnight incubation under static conditions facilitated the formation of a uniform array of stem cell aggregates within individual chambers. Subsequent to this, the organoids were perfused with mTeSR1 for one day before initiating a 40-day-long differentiation program under continuous perfusion conditions. Media was deposited to the reservoirs, which have open tops allowing for gas equilibrium between the media and the atmosphere inside a culture incubator. Additionally, the high gas permeability of PDMS supports the enhanced diffusion of CO_2_ and oxygen to the cells (e.g., D_PDMS_ = 3.2×10^−9^ m^2^/s vs. D_water_ = 2.3×10^−9^ m^2^/s^66,67^)

Our differentiation approach, referring to an earlier study^68^, was structured into three distinct phases: progenitor induction, primary maturation of dopaminergic neurons, and advanced maturation of neuronal networks. Each day, a freshly prepared basal media (DMEM/F12 supplemented with 1% (v/v) Glutamax, 0.5% (v/v) N2 supplement, 1% (v/v) B27 supplement without vitamin A) with differentiation factors were deposited into the media reservoirs. The perfusion rate was designed at 150 µL per day per ten organoids during the progenitor induction phase, transitioning to 300 µL per day per ten organoids thereafter. This perfusion configuration was designed to establish a flow through two parallel sets of five serial organoids.

In the early stages of differentiation (D1-D12), the transition from pluripotent state to midbrain floor-plate precursors was initiated by applying small molecules. Specifically, LDN193189 (100 nM; D1-12; STEMCELL Technologies) and SB431542 (10 µM; D1-8; STEMCELL Technologies) were utilized to inhibit BMP and TGF-β signaling, respectively. Furthermore, Purmorphamine (2 µM, D3-8; Tocris) and SAG (1 µM, D3-8; STEMCELL Technologies) were introduced to activate Sonic Hedgehog (SHH) signaling. During later precursor patterning, the addition of the WNT signaling activator, a low concentration of CHIR99021 (1.5 µM; D5-12; Tocris), was used. Upon reaching the maturation stage from D13 onwards, basal components included neurotrophic factors (BDNF and GDNF; 10 ng/ml; PeproTech) and Ascorbic acid (0.2 mM). The initiation of maturation involved the application of DAPT (N-[N-(3,5-difluorophenacetyl)-L-alanyl]-S-phenylglycine t-butyl ester; 10 µM; D13-36; Tocris) as a notch signaling inhibitor, alongside dcAMP (dibutyryl cyclic AMP; 100 µM; D13-36; Tocris) for cAMP/PKA/CREB activation. A transition from DMEM/F12 to BrainPhys™ Neuronal Media (STEMCELL Technologies) was gradually implemented between D37 and D40 to promote neuronal activities while maintaining the trophic factors and Ascorbic acid.

### MEA Electrophysiology of Midbrain Organoids

Midbrain organoids differentiating from pluripotent stem cells to floor-plate progenitors were retrieved on day 15 of MAP differentiation and placed on Laminin/Poly-L-Ornithine (PLO)-coated MEA substrates, fabricated using the same method as the MAP electrode. The remaining differentiation protocol was carried out in a static MEA culture condition, with partial medium refreshment every 8 hours, replacing a quarter of the volume to maintain compatibility with MAP’s biochemical conditioning. To assess the impact of this medium refreshment, we also transferred a set of midbrain organoids to round-bottom 96-well plates and cultured them using the same medium refreshment protocol, which resulted in developmental characteristics compatible with those of MAP organoids (Supplementary Fig. 3). The figure also shows that the integration of electrodes in the MAP caused no developmental alterations. L-DOPA was introduced by adding 5 mM to the fresh medium, resulting in a final concentration of 50 µM in the MEA culture. MPP^+^ was applied by initially adding it at a 1 mM concentration to achieve a 10 µM final concentration in the MEA culture, followed by regular medium refreshment with 10 µM MPP^+^ in a quarter volume. For clarity in microscopy imaging, plain non-electrode glass substrates with the same SiO_2_ deposition and Laminin/PLO coating were used. Electrophysiological recordings were taken from electrodes positioned beneath organoids, selected via visual examination, paired with a distant reference electrode in a Faraday cage. External stimulation to characterize the readout of non-contact signaling was performed by positioning a pair of syringe needles, separated by 1 mm, 1 mm above a target electrode, and applying a 15 Hz, 20 µV sinusoidal oscillation using a function generator.

### Generation of Cortical Organoids

The human stem cell-derived cortical brain organoids were prepared following the published literature^69^. Briefly, HES3 hESCs, cultured on Matrigel-coated dishes, were seeded into wells of U-bottom ultra-low-attachment 96-well plate containing neural induction medium. This medium comprised DMEM-F12, 15% (v/v) KSR, 5% (v/v) heat-inactivated FBS (Life Technologies), 1% (v/v) Glutamax, 1% (v/v) MEM-NEAA, and 100 µM β-Mercaptoethanol, supplemented with 10 µM SB-431542, 100 nM LDN-193189, 2 µM XAV-939, and 50 µM Y27632. On day 2 and day 4, FBS and Y27632 were sequentially removed. This medium was refreshed every alternate day until day 10 when the organoids were transferred to an ultra-low-attachment six-well plate.

The organoids were nurtured in a spinning medium without vitamin A for the subsequent stages. This medium was a 1:1 mixture of DMEM-F12 and Neurobasal media, containing 0.5% (v/v) N2 supplement, 1% (v/v) B27 supplement without vitamin A, 0.5% (v/v) MEM-NEAA, 1% (v/v) Glutamax, 50 µM β-Mercaptoethanol, and 0.025% Insulin. This medium was replaced every other day until day 18, at which point the medium was transitioned to the medium with additional vitamin A, using B27 supplement with vitamin A. This medium was supplemented with 20 ng/ml BDNF and 200 µM ascorbic acid. Following day 18, the medium exchange occurred every 4 days.

From day 35 onwards, the organoids were transferred to the integrated MAP device for concurrent EEG recording and perfusion.

### Immunostaining and Transcriptomic Analysis

Immunostaining was conducted using MAP perfusion, encompassing the following general steps through perfusion: (1) 8-hour perfusion of 4% paraformaldehyde (PFA) in PBS. (2) 8-hour perfusion of PBS for thorough washing. (3) 8-hour perfusion of 0.1% triton-X for permeabilization. (4) 1-day perfusion of 2% BSA in PBST for blocking. (5) 2-day perfusion of primary antibodies in 1% BSA in PBST. (6) 1-day perfusion in 1% BSA in PBST. (7) 2-day perfusion of secondary antibodies in 1% BSA in PBST. (8) 1-day perfusion of PBST and PBS. (9) 8-hour perfusion of an optical clearing solution (RapiClear 1.52; SunJin Lab).

The antibodies used were anti-MAP2 (822501, BioLegend), anti-TH (ab137869, Abcam), anti-S100B (S2532, MilliporeSigma), and anti-Tuji1(A-21435, ThermoFisher Scientific) diluted at 1:10000, 1:300, 1:1000, and 1:1000 respectively. Secondary antibodies were used at a 1:800 dilution to match the target species (ThermoFisher Scientific). Immunostaining was visualized with an inverted laser scanning confocal microscope (Zeiss LMS980), while the organoid remained positioned within the device. Z-stacks with a thickness of 10 µm were generated using ImageJ (ver 1.54; National Institutes of Health). MAP2- and GFAP-positive regions were quantified within 50 × 50 µm² areas, with thresholds defined by antibody controls.

For RNA extraction, RNeasy Plus Mini Kit (Qiagen) was used. The process involved perfusing a lysis buffer (Buffer RLT Plus) through the organoid chambers for on-chip lysis, followed by retrieval of lysates from the chip for pooled RNA extraction. For single-organoid analysis, individual organoids were retrieved by carefully opening the device with a blade and lysed in microcentrifuge tubes using Buffer RLT Plus. Following the manufacturer’s protocol, RNA was collected and immediately converted into complementary DNA (cDNA) using a reverse transcription kit (iScript Reverse Transcription Supermix, Bio-Rad). The resulting cDNA samples were stored at −80°C before PCR analysis. Three biological replicates were prepared from separate experiments for statistical analysis.

Real-time PCR analysis was performed using the SsoAdvanced Universal SYBR Green Supermix (Bio-Rad) and the CFX96 real-time PCR system (Bio-Rad). The PCR cycle consisted of 40 cycles with denaturation at 95°C and annealing at 60°C. PCR byproduct amplicons were verified through melting analysis (65-95°C in 0.5°C increments). A list of primers used is available in the Supplementary Table 1.

To examine the developmental trajectory in Fig. 2, 12 genes related to midbrain organoid development except for pluripotency and progenitor from the Fig. 3d list were quantified from the single-organoid RNA extractions. To interpret the overall gene variation during differentiation, we conducted a principal component analysis of the z-score normalized 12-gene expression at five time points, including 7, 14, and 28 days after plating on differentiation day 15.

### EEG Data Analysis

EEG data, recorded at a 5 kHz sampling rate, underwent analysis for local field potential, spiking activity, brainwaves, and power spectrum, using a MATLAB code (Supplementary Fig. 6). The initial step involved notch-filtering the raw data to eliminate environmental noise at 60 Hz (±3 Hz). Next, the data was transformed into local field potential by applying a 100 Hz lowpass filter and into spike activity via a 300 Hz highpass filter, then further converted to entire unit activity (EUA) by rectification. For brainwave analysis, the local field potential was segregated into distinct frequency bands: delta (0.5-4 Hz), theta (4-8 Hz), alpha (8-12 Hz), beta (12-35 Hz), and gamma (35-100 Hz), accomplished through corresponding bandpass filtering. These filters were designed as infinite impulse response filters.

To generate a spectrogram, the raw data underwent short-time Fast Fourier Transform (FFT) analysis on 20-second-long segments with a resolution of 2 seconds. Also, wavelet analysis was used to obtain a better temporal resolution in spectrogram analysis using a continuous 1-dimensional wavelet transform (CWT) with a Morse wavelet and a VoicesPerOctave set at 16. Further details of the analytical process can be found in Supplementary Fig. 6.

For phase-amplitude coupling (PAC), the phase of the slower oscillatory signal was extracted in the range of 0.5-3.5Hz with 1Hz bandwidth in a resolution of 1Hz. For the wave extraction, finite impulse response (FIR) filters were used for its accuracy. Then, the phase was extracted from the complex domain of Hilbert transform. Similarly, the amplitude of the fast oscillation signal was calculated using the serial application of FIR filtering and the amplitude calculation of Hilbert transformed complex signal. For the amplitude, the range of 10-60Hz was analyzed with a 2 Hz bandwidth in a resolution of 1 Hz. The phase values were divided into bins with a width of 20°. To calculate the modulation index, we adopted a method to calculate the Kullback–Leibler (KL) distance from a uniform distribution describing no phase-amplitude coupling^70^. The time-resolved modulation index was quantified using 1-minute segments of raw data, with a resolution of 0.5 minutes.

### Comparison of Organoid Culture Methods

We compared the presented MAP culture with other alternatives to highlight the advancements MAP offers in cultivating brain organoids. For this comparison, we designed an overflow fluidic device and a bioreactor culture. The overflow fluidic device shared the overall dimensions of the MAP system but lacked the nanochannel separation between organoid chambers and perfusion channels. For the bioreactor culture, we first generated stem cell aggregates using Aggrewell 800 (STEMCELL Technologies) according to the manufacturer’s online protocol, then transferred them to a magnetic stirring flask spinning at 65 rpm (Corning Glass Spinner Flask). The media refreshment of the overflow culture was designed to use the same media reservoirs and tilting rate as the MAP system, while the bioreactor culture followed the recommended protocol^71^. Glucose and lactate concentrations were quantified from regularly collected samples using commercial kits, following the manufacturer protocols (Glucose & Lactate Assay Kit; Abcam).

### Statistical Analysis

All statistical tests performed are described in the figure legends. Sample sizes are provided in the figure legends. A two-tailed unpaired Student’s t-test assuming unequal variances was conducted to statistically analyze differences between two groups. Repeated measures ANOVA followed by Tukey’s Honestly Significant Difference (HSD) post-hoc analysis was applied to assess differences over multiple time points. The 10% chosen in Fig. 4f was selected to balance between focusing on notable patterns in PAC changes associated with LFP amplitude while capturing enough data to avoid the noise inherent in the extremes. Microsoft Excel was used to perform statistical analysis. All p values are displayed in the figure legends.

