## Supplementary Information for "Brainwaves Monitoring via Human Midbrain Organoids Microphysiological Analysis Platform: MAP"

### Contents

- *Supplementary Figure 1-9*
- *Supplementary Table 1*

**Supplementary Figure 1. Simulation of MAP Fluidic Unit**

The MAP fluidic unit, comprising two perfusion channels, a hemispherical chamber, and an array of nanochannel perfusion barriers, was simulated using COMSOL Multiphysics (ver. 5.2). The dimensions are found in Extended Data Fig. 1. **a,** The sectional profile of flow velocity at the middle height of the perfusion channel is shown in the top image, and the AA' section is displayed in the bottom image. **b,** The MAP fluidic unit, comprising two perfusion channels, a hemispherical chamber and an array of nanochannel perfusion barriers, was simulated using COMSOL Multiphysics (ver. 5.2). **c,** Another fluidic design without the nanochannel perfusion barriers was simulated to assess the endothelium-like barrier's role in regulating the fluidic shear stress in the hemispherical chamber. Spatial profiles of fluidic shear stress were computed on the surfaces of pseudo-organoids modeled with diameters of 50, 100, 200, 500, and 1000 μm. It should be noted that the two sets are displayed using different colormap scales.

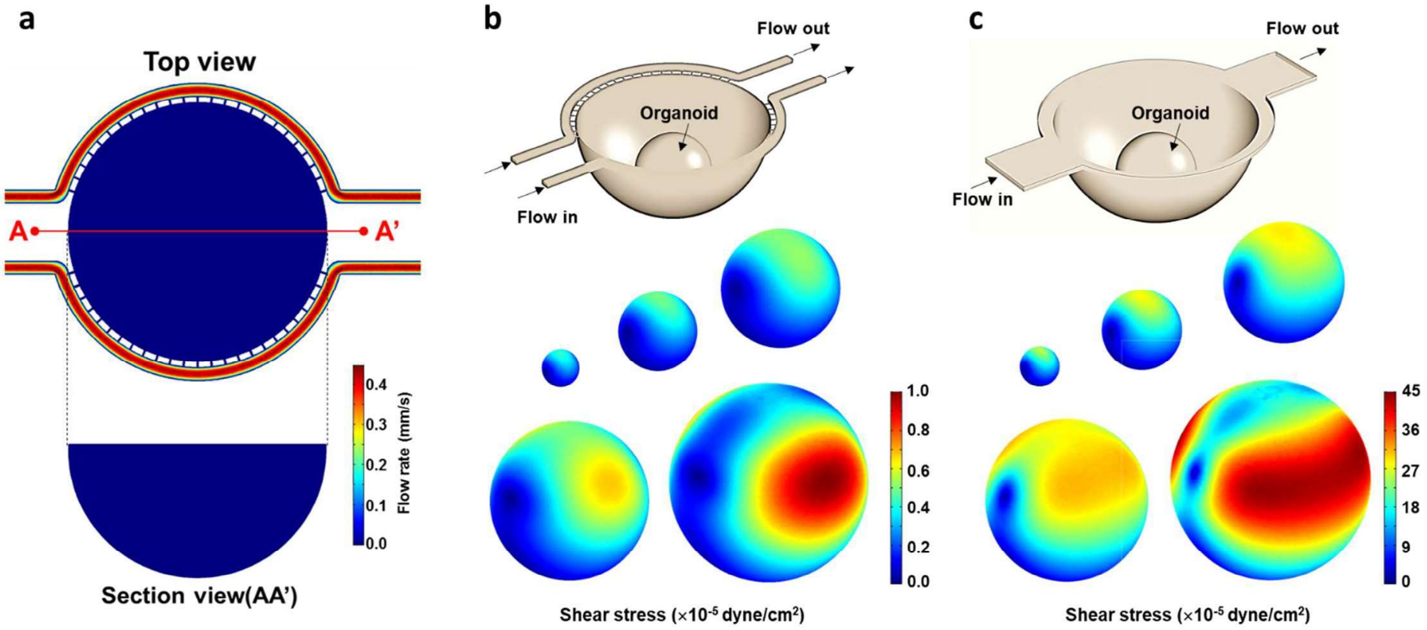


**Supplementary Figure 2. Midbrain Organoid Development in MAP**

**a,** Optical tracing of midbrain organoid growth in MAP. The progression of midbrain organoid development within the MAP system is illustrated through an optical trace. Through the timely applied differentiation factors (details in Methods), a minute aggregate of stem cells is adeptly nurtured into spherical midbrain constructs. **b,** Growth pattern of midbrain organoids. The bar chart portrays the growth pattern of midbrain organoids within the MAP platform. The bar height presents the mean value, accompanied by error bars indicating the standard deviation. This characterization was derived from data from 20 organoids. The left circle is a representative equivalent circle, facilitating visualization. **c,** Analysis of midbrain organoid morphological characteristics. Normalized perimeter and circularity were analyzed for midbrain organoids at day 80, which developed from two initial cell numbers in the experimental protocol. Each organoid's data is represented as dots on the graph, with error bars indicating standard deviation (n=20). **d,** Precision in organoid modeling in MAP with initial stem cell numbers. Photographs of midbrain organoid arrays captured on day 80 show the uniformity and precision achieved in organoid generation using the MAP system. The images affirm the platform's capability for controlled and consistent organoid development. In both panels (Scale bar: 200 μm).

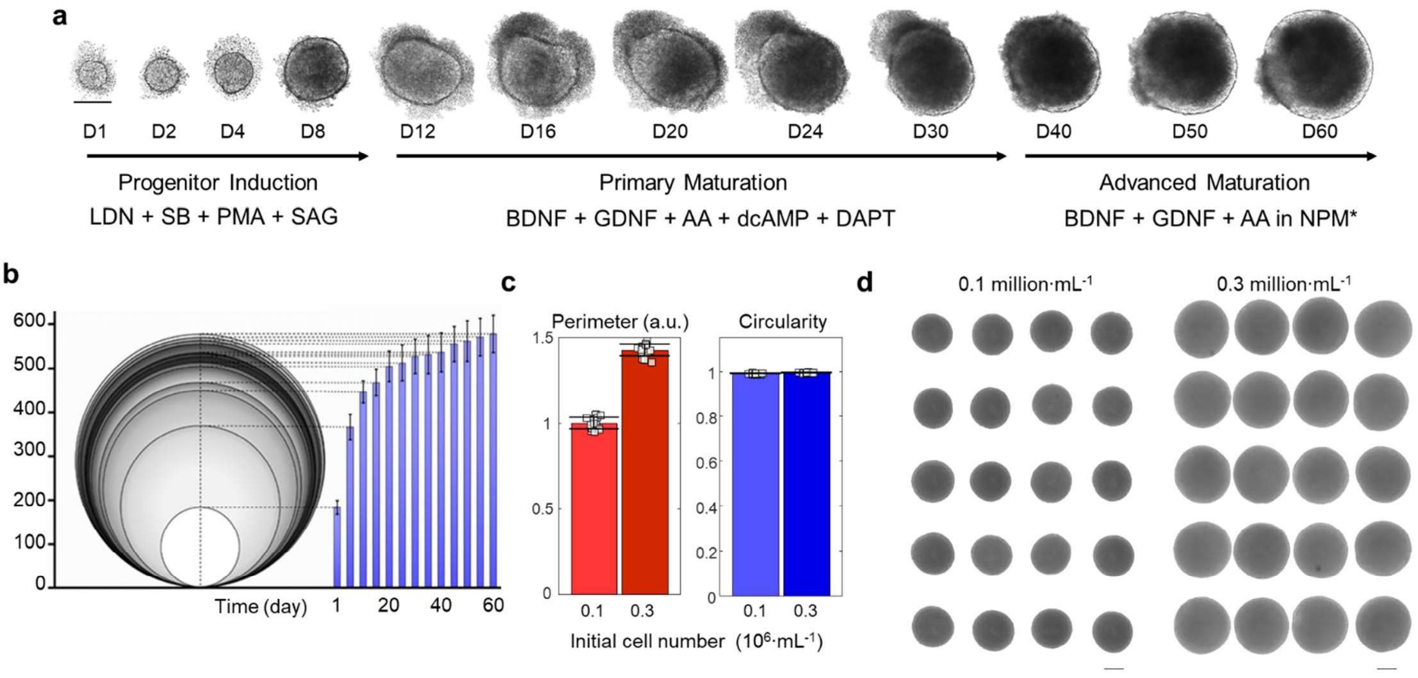


**Supplementary Figure 3. Gene Expression Comparison among Midbrain Organoids in MAP, MEA, and Multiwell Cultures.**

Fig. S3a demonstrates that the media refreshment protocol described in the main text and Methods, designed for our presented MEA electrophysiology study, allows for a valid gene expression comparison between organoids developed in MAP and MEA cultures. The MEA media conditioning, involving an 8-hour interval with a quarter-volume media refreshment, was implemented to ensure biochemical similarity between MAP and MEA cultures. To exclude the impact of the fluid dynamic difference on gene expression changes, we generated an additional set of organoids in round-bottom, non-adherent 96-well plates with the same media refreshment method as the MEA culture. The gene expression results from single organoids are overlaid on Fig. 2d, showing a compatible developmental trajectory between midbrain organoids from the 96-well plates and those from MAP culture. While MAP organoids exhibit greater homogeneity among individual organoids compared to the others, the results suggest that differences in media refreshment have a negligible effect on gene expression across the models.

While the MEA-plated development showed significant deviations in the developmental trajectory, the integration of electrodes within the MAP resulted in an insignificant effect on midbrain organoid development, as shown in Supplementary Fig. 3b.

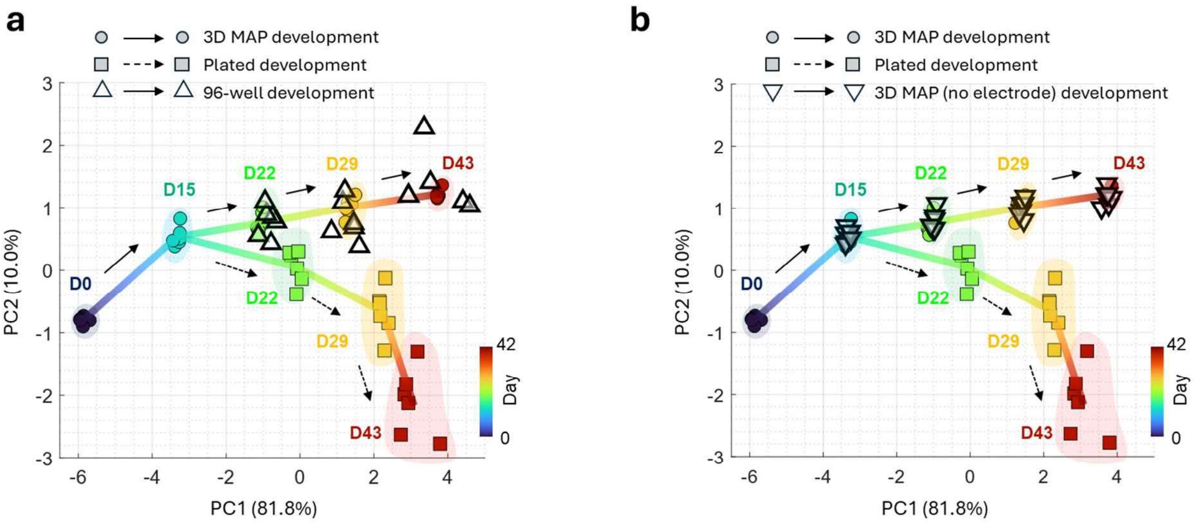


**Supplementary Figure 4. Simulation models of MEA and MAP electrophysiological setups.**

**a,** The contact configuration of the MEA readout on contacting cells can be represented by the equivalent circuit shown in the left part of (i), where $R_{seal}$ (a resistance generated by the tight adhesion of the cell membrane to an electrode) is designed to be high to enhance sensitivity to the activity of contacting cells ($V_A$). However, this configuration also introduces a high and temporally variant resistance in the signal pathway connecting distant cell activity ($V_B$), posing challenges in accurately capturing distant electrophysiological activities. On the other hand, the MAP's configuration prevents direct contact between the cells and the electrode by maintaining a distance between the organoid and the electrodes. Although this distance may limit the detection of single-cell activities, it allows for the highlighting of synchronized activities among electrogenic cells. This setup is analogous to how EEG analyzes tissue-level activities, capturing population-averaged dipole-like signals as shown in (ii). **b**, The enclosed boundary of the MAP's hemispherical chamber significantly enhances the sensitivity of electrophysiological readouts by preventing the dissipation of electric potential that typically occurs under open boundary conditions. Computational simulations illustrate the electric potential distributions around a neuron-like dipole oscillation positioned at the same relative distance from an electrode. These simulations demonstrate that the potential can be amplified by 250% when the hemispherical chamber boundary is integrated. In the simulation, the conductivities of the media, glass, and MAP chamber were estimated to be 1.66 S/m, $1\times10^{-11}$ S/m, and $1\times10^{-12}$ S/m, respectively.

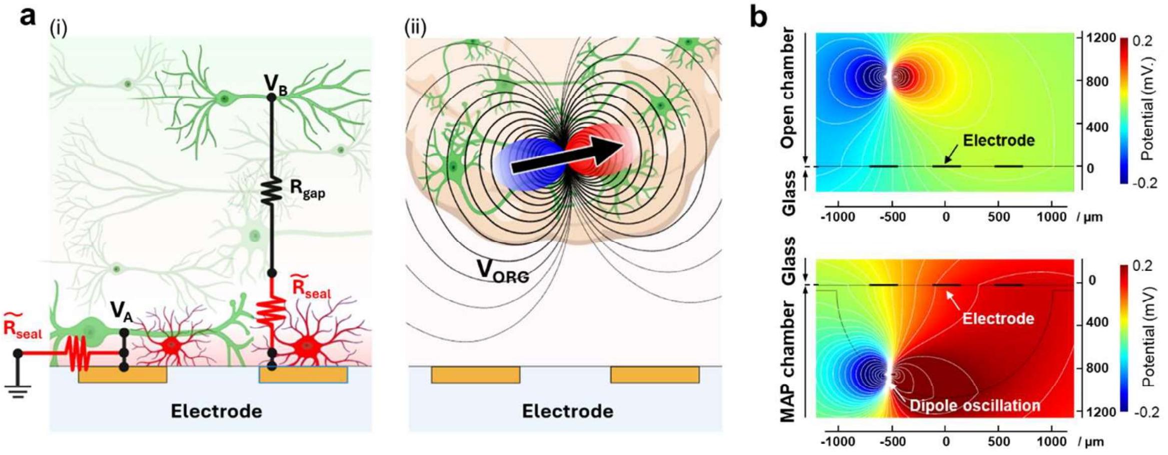


**Supplementary Figure 5. Calcium imaging of midbrain organoids using live-cell cytosolic calcium dye.**

**a,** Pseudo-colored fluorescent image on a midbrain organoid at day 68. The imaging was focused on the organoid's middle height using epifluorescent microscopy. The color map transitions from blue to red, representing a gradient from low to high fluorescent signals of Rhod-3. **b,** Fluorescence intensity time traces for regions of interest (ROIs) indicated in panel a. The dot red lines highlight time-correlated events among these ROIs. **c,** Calcium propagation profile in a close-up profile. This illustration highlights that a localized initiation of cytoplasmic calcium increase triggers the spatial propagation of calcium fluctuation, exemplifying sub-second dynamics. **d,** Pseudo-colored image of calcium dye (Fluo-4) on a midbrain organoid at day 65. The focal plane was on the organoid’s middle height using epifluorescent microscopy. For the correlation analysis, the image recording was extended to 30 sec. **e,** Fluorescence intensity traces on regions of interest (ROIs) indicated in panel d. **f,** Calcium signal correlation analysis against the annotated local points conducted using MATLAB's pdist function with the 'correlation' option. This approach enabled the comparison of spatial synchrony in individual pixels of the calcium fluorescence movie, revealing the presence of multiple local networks within the midbrain organoids.

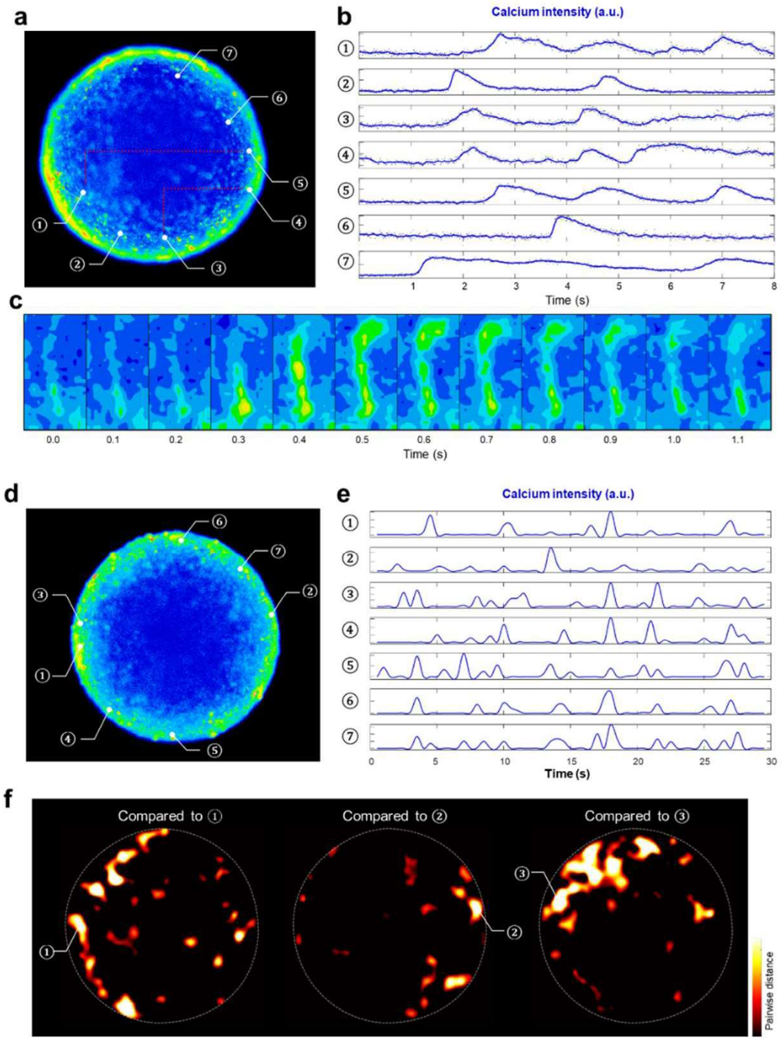


**Supplementary Figure 6. Post-hoc Signal Processing of Brain Organoid EEG from MAP**. This figure presents an overview of the post-hoc signal processing pipeline applied to the brain organoid electroencephalogram (EEG) data acquired within MAP. The raw signals recorded from each electrode were sampled at a rate of 5 kHz. Subsequently, a custom MATLAB code was employed to perform a sequential processing sequence, deriving multiple parameters and features. The first step was to derive EEG signals by filtering out environmental noise at 60Hz and LFP (local field potential) signals. In this study's data, EEG and LFP are depicted as overlaid black and red lines. Physiologically, LPF offers insights into the collective neural activity within the organoids. Then, brainwaves were extracted from the LFP signal, spanning the frequency ranges of 0.5-4 Hz for delta, 4-8 Hz for theta, 8-12 Hz for alpha, 12-35 Hz for beta, and 35-100 Hz for gamma rhythms. These brainwave components provide information about different neural states and functions. In addition, spiking activity above 300 Hz was detected within the EEG-like signals, providing an understanding of the high-frequency neural firing of individual neurons. The power density spectra, which reveal the distribution of signal power across different frequencies, were calculated further to analyze the frequency composition of the EEG-like signals. The systematic application of this signal processing methodology led to the transformation of the initially acquired EEG-like signals into an extensive array of interpretable parameters and features. These outcomes are relatable within conventional physiological contexts, enhancing the understanding of the intricate neural dynamics within the brain organoids studied.

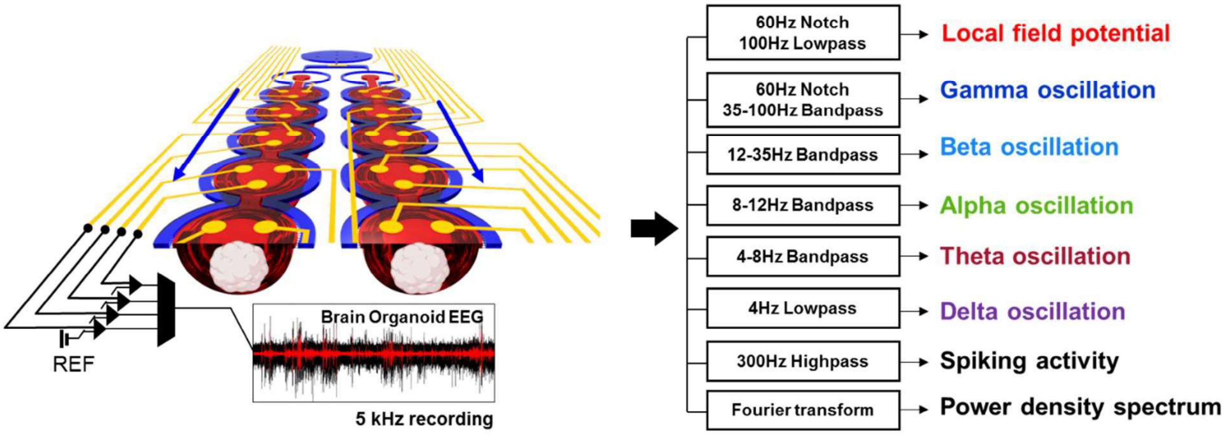


**Supplementary Figure 7. TH mRNA change induced by 10 μM MPP+ application.**

The expression levels were quantified by comparing them with age-matched vehicle controls in RT-qPCR analysis of RNA samples from 10 organoids. The dotted line represents the average, and the shaded area depicts the standard deviation of biological replicates (n=3). The repeated measures ANOVA followed by Tukey's HSD test revealed statistically significant decreases in TH expression after day 15. **($p < 0.01$) and ***($p < 0.001$).

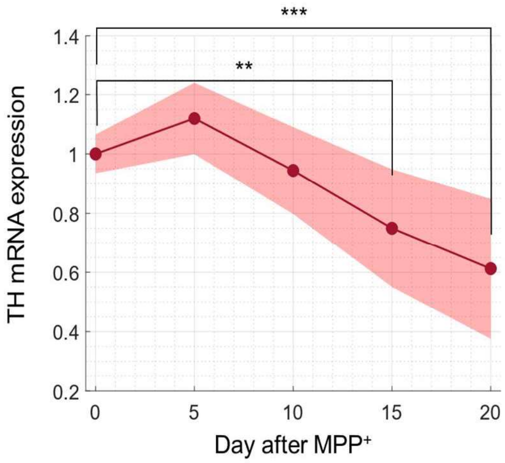

**Supplementary Figure 8. Characterization of control and proximal readouts in MAP. a,** Temporal changes of a control electrode without organoids compared to representative readouts per case. The controlled experimental environment and consistent fluidics produce a low-level blank signal of less than 35 μVpp. **b,** Correlation analysis among electrodes in MAP. (i) Three-channel positions above an organoid named "D" in MAP. (ii) Electrical readouts of three channels on organoid "D", a channel on organoid "C" in an upstream chamber, and a channel on organoid "X" in a distant chamber connected to a separate perfusion unit. The three channels (D①, D②, D③) exhibit well-synchronized temporal patterns with localized variations attributed to relative changes in source geometries within the organoid. Electrodes not assigned to the same organoids display poor synchronization, reflecting different organoid activities. (iii) Signal correlation of ten organoids within a perfusion unit. The heatmap summarizing correlation coefficients for five-minute readouts on ten individual organoids presents insignificance of signal synchrony as addressing different organoids. Note that the heatmap color scale has a break between 0.014 and 0.998.

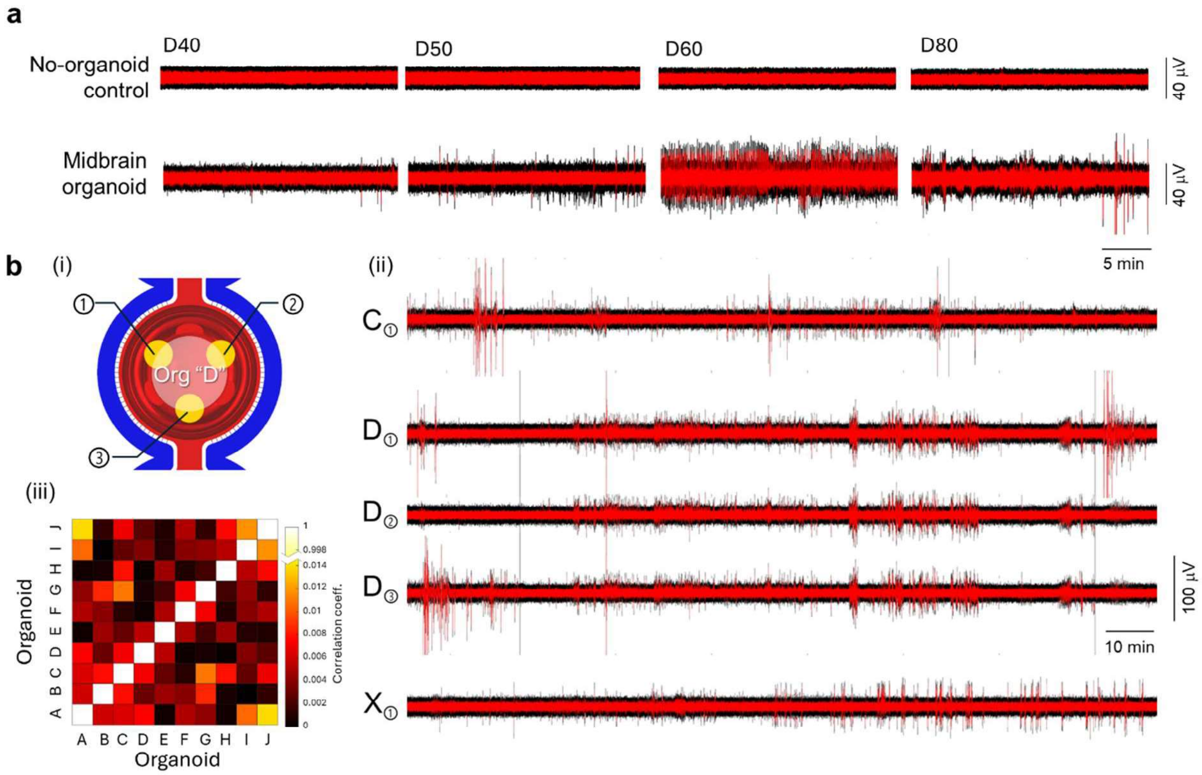

**Supplementary Figure 9. Midbrain organoid brainwave traces during 10 µM⁺ MPP perfusion in MAP.** The time trace of MAP's recording, displayed with delta (purple), theta (red), alpha (green), beta (light blue), and gamma (dark blue) brainwave rhythms, illustrates global transitions in brainwave activity from hyperactive to inactive states as shown in Figure 4. Meanwhile, localized events demonstrate intriguing characteristics that merit advanced analysis in future studies.

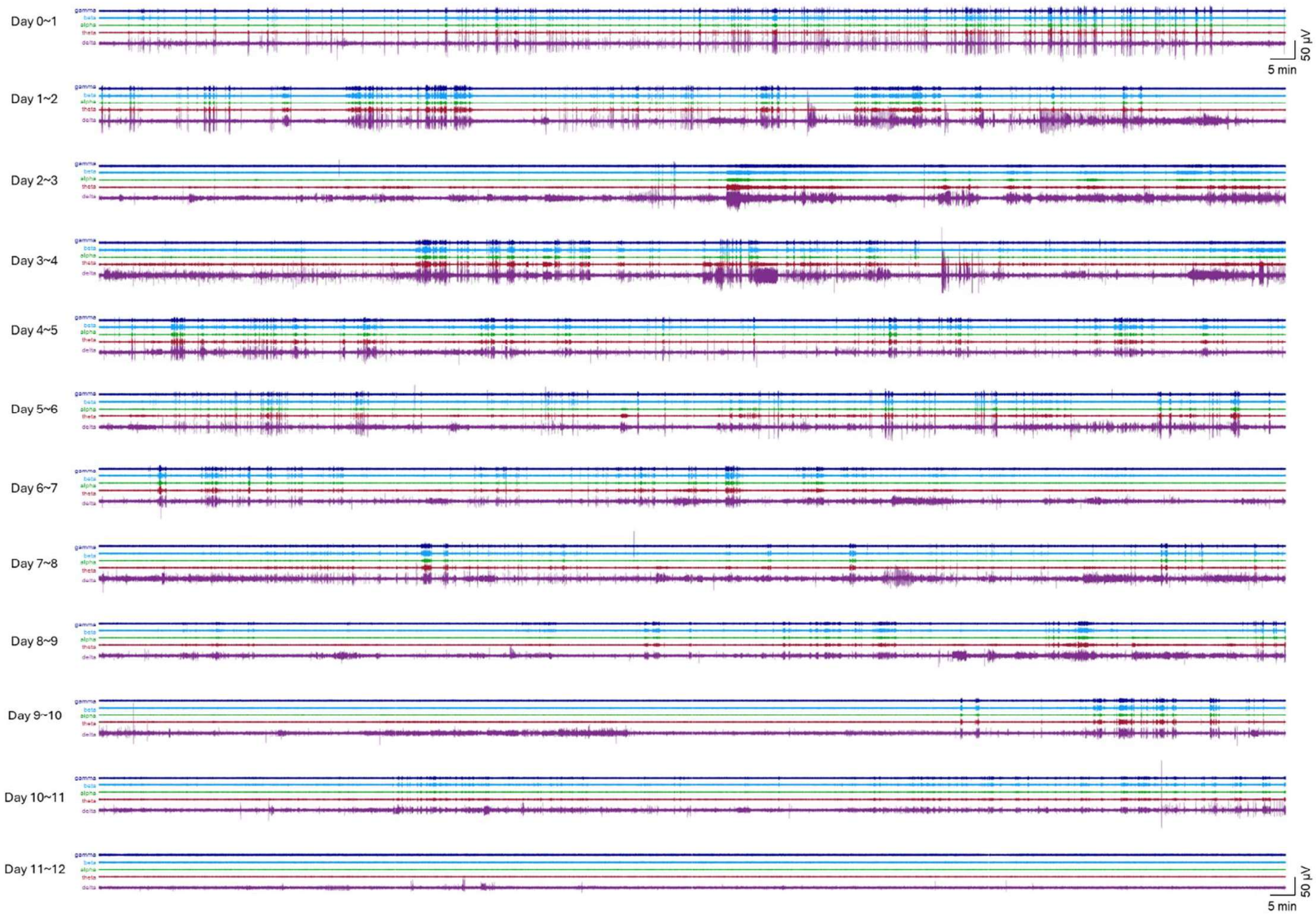
